# The conserved *PFT1* tandem repeat is crucial for proper flowering in *Arabidopsis thaliana*

**DOI:** 10.1101/006437

**Authors:** Pauline Rival, Maximilian O. Press, Jacob Bale, Tanya Grancharova, Soledad F. Undurraga, Christine Queitsch

**Affiliations:** University of Washington Department of Genome Sciences; University of Washington Molecular and Cellular Biology Graduate Program; University of Washington Department of Biochemistry; current address: Universidad Mayor Centro de Genómica y Bioinformática, Santiago, Chile

**Keywords:** PFT1, MED25, short tandem repeat, flowering, microsatellite

## Abstract

It is widely appreciated that short tandem repeat (STR) variation underlies substantial phenotypic variation in organisms. Some propose that the high mutation rates of STRs in functional genomic regions facilitate evolutionary adaptation. Despite their high mutation rate, some STRs show little to no variation in populations. One such STR occurs in the *Arabidopsis thaliana* gene *PFT1* (*MED25*), where it encodes an interrupted polyglutamine tract. Though the *PFT1* STR is large (∼270 bp), and thus expected to be extremely variable, it shows only minuscule variation across *A. thaliana* strains. We hypothesized that the *PFT1* STR is under selective constraint, due to previously undescribed roles in PFT1 function. We investigated this hypothesis using plants expressing transgenic *PFT1* constructs with either an endogenous STR or with synthetic STRs of varying length. Transgenic plants carrying the endogenous *PFT1* STR generally performed best across adult PFT1-dependent traits, in terms of complementing a *pft1* null mutant. In stark contrast, transgenic plants carrying a *PFT1* transgene lacking the STR entirely phenocopied a *pft1* loss-of-function mutant for flowering time phenotypes, and were generally hypomorphic for other traits, establishing the functional importance of this domain. Transgenic plants carrying various synthetic constructs occupied the phenotypic space between wild-type and *pft1*-loss-of-function mutants. By varying *PFT1* STR length, we discovered that *PFT1* can act as either an activator or repressor of flowering in a photoperiod-dependent manner. We conclude that the *PFT1* STR is constrained to its approximate wild-type length by its various functional requirements. Our study implies that there is strong selection on STRs not only to generate allelic diversity, but also to maintain certain lengths pursuant to optimal molecular function.

## INTRODUCTION

Short tandem repeats (STRs, microsatellites) are ubiquitous and unstable genomic elements that have extremely high mutation rates (Subramanian *et al*. 2003; Legendre *et al*. 2007; Eckert and Hile 2009), leading to STR copy number variation within populations. STR variation in coding and regulatory regions can have significant phenotypic consequences (Gemayel *et al*. 2010). For example, several devastating human diseases, including Huntington’s disease and spinocerebellar ataxias, are caused by expanded STR alleles (Hannan 2010). However, STR variation can also confer beneficial phenotypic variation and may facilitate adaptation to new environments (Fondon *et al*. 2008; Gemayel *et al*. 2010). For example, in *Saccharomyces cerevisiae* natural polyQ variation in the FLO1 protein underlies variation in flocculation, which is important for stress resistance and biofilm formation in yeasts (Verstrepen *et al*. 2005). Natural STR variants of the *Arabidopsis thaliana* gene *ELF3*, which encode variable polyQ tracts, can phenocopy *elf3* loss-of-function phenotypes in a common reference background (Undurraga *et al*. 2012). Moreover, the phenotypic effects of *ELF3* STR variants differed dramatically between the divergent backgrounds Col and Ws, consistent with the existence of background-specific modifiers. Genetic incompatibilities involving variation in several other STRs have been described in plants, flies, and fish (Peixoto *et al*. 1998; Scarpino *et al*. 2013; Rosas *et al*. 2014). Taken together, these observations argue that STR variation underlies substantial phenotypic variation, and may also underlie some genetic incompatibilities.

The *A. thaliana* gene *PHYTOCHROME AND FLOWERING TIME 1* (*PFT1, MEDIATOR 25, MED25*) contains an STR of unknown function. In contrast to the comparatively short and pure *ELF3* STR, the *PFT1* STR encodes a long (∼90 amino acids in PFT1, vs. 7-29 for ELF3), periodically interrupted polyQ tract. The far greater length of the *PFT1* STR leads to the prediction that its allelic variation should be greater than that of the highly variable *ELF3* STR (Legendre *et al*. 2007, http://www.igs.cnrs-mrs.fr/TandemRepeat/Plant/index.php). However, in a set of diverse *A. thaliana* strains, *PFT1* STR variation was negligible compared to that of the *ELF3* STR (Supp. Table 1). Also, unlike ELF3, the PFT1 polyQ is conserved in plants as distant as rice, though its purity decreases with increasing evolutionary distance from *A. thaliana*. A glutamine-rich C-terminus is conserved even in metazoan MED25 (File S1). Recent studies of coding STRs suggested that there may be different classes of STR. Specifically, tandem repeats that are conserved across large evolutionary distances appear in genes with substantially different functions than those coding tandem repeats that are not strongly conserved in species (Schaper *et al*. 2014). Consequently, *PFT1/MED25* polyQ conservation may functionally differentiate the *PFT1* STR from the *ELF3* STR.

*PFT1* encodes a subunit of Mediator, a conserved multi-subunit complex that acts as a molecular bridge between enhancer-bound transcriptional regulators and RNA polymerase II to initiate transcription (Bäckström *et al*. 2007; Conaway and Conaway 2011). *PFT1*/*MED25* is shared across multicellular organisms but absent in yeast. In *A. thaliana*, the PFT1 protein binds to at least 19 different transcription factors (Elfving *et al*. 2011; Ou *et al*. 2011; Çevik *et al*. 2012; Chen *et al*. 2012) and has known roles in regulating a diverse set of processes such as organ size determination (Xu and Li 2011), ROS signaling in roots (Sundaravelpandian *et al*. 2013), biotic and abiotic stress (Elfving *et al*. 2011; Kidd *et al*. 2009; Chen *et al*. 2012), phyB-mediated-light signaling, shade avoidance and flowering (Cerdán and Chory 2003; Wollenberg *et al*. 2008; Iñigo, Alvarez, *et al*. 2012; Klose *et al*. 2012).

PFT1 was initially identified as a nuclear protein that negatively regulates the phyB pathway to promote flowering in response to specific light conditions (Cerdán and Chory 2003; Wollenberg *et al*. 2008). Recently, Iñigo and colleagues (2012) showed that PFT1 activates *CONSTANS* (*CO*) transcription and *FLOWERING LOCUS T* (*FT*) transcription in a *CO*-independent manner. Specifically, proteasome-dependent degradation of PFT1 is required to activate *FT* transcription and to promote flowering (Iñigo, Giraldez, *et al*. 2012). The wide range of PFT1-dependent phenotypes is unsurprising given its function in transcription initiation, yet it remains poorly understood how PFT1 integrates these many signaling pathways.

Given the conservation of the PFT1 polyQ tract and the known propensity of polyQ tracts for protein-protein and protein-DNA interactions (Escher *et al*. 2000; Schaefer *et al*. 2012), we hypothesized that this polyQ tract plays a role in the integration of multiple signaling pathways and is hence functionally constrained in length. We tested this hypothesis by generating transgenic lines expressing *PFT1* with STRs of variable length and evaluating these lines for several PFT1-dependent developmental phenotypes. We show that the *PFT1* STR is crucial for *PFT1* function, and that PFT1-dependent phenotypes vary significantly with the length of the *PFT1* STR. Specifically, the endogenous STR allele performed best for complementing the flowering and shade avoidance defects of the *pft1-2* null mutant, though not for early seedling phenotypes. Our data indicate that most assayed *PFT1*-dependent phenotypes require a permissive *PFT1* STR length. Taken together, our results suggest that the natural *PFT1* STR length is constrained by the requirement of integrating multiple signaling pathways to determine diverse adult phenotypes.

## RESULTS

We used amplification fragment length polymorphism analysis and Sanger sequencing to evaluate our expectation of high *PFT1* STR variation across *A. thaliana* strains. However, we observed only three alleles of very similar size (encoding 88, 89 and 90 amino acids, Table S1), in contrast to six different alleles of the much shorter *ELF3 STR* among these strains, some of which are three times the length of the reference allele (Undurraga *et al*. 2012). These data implied that the *PFT1* and *ELF3* STRs respond to different selective pressures. In coding STRs, high variation has been associated with positive selection (Laidlaw *et al*. 2007), though some basal level of neutral variation is expected due to the high mutation rate of STRs. We hypothesized that the *PFT1* STR was constrained to this particular length by *PFT1*’s functional requirements.

To test this hypothesis, we generated transgenic *A. thaliana* carrying *PFT1* transgenes with various STR lengths in an isogenic *pft1*-2 mutant background. These transgenics included an empty vector control (VC), 0R, 0.34R, 0.5R, .75R, 1R (endogenous *PFT1* STR allele), 1.27R, and 1.5R constructs. All STRs are given as their approximate proportion of WT STR length – for instance, the 1R transgenic line contains the WT STR allele in the *pft1*-2 background (Table S2). We used expression analysis to select transgenic lines with similar *PFT1* expression levels (Table S3).

### The *PFT1* STR length is essential for wild-type flowering and shade avoidance

We first evaluated the functionality of the different transgenic lines in flowering phenotypes. Removing the STR entirely substantially delayed flowering under long days (LD, phenotypes days to flower, rosette leaf number at flowering; Figure 1A). In LD, any STR allele other than 0R was able to rescue the *pft1-2* late-flowering phenotype. Indeed, one allele (1.5R) showed earlier flowering than WT (Figure 1B, 1C), whereas other alleles provided a complete or nearly complete rescue of the *pft1-2* mutant (Figure 1D).

**Figure 1.**
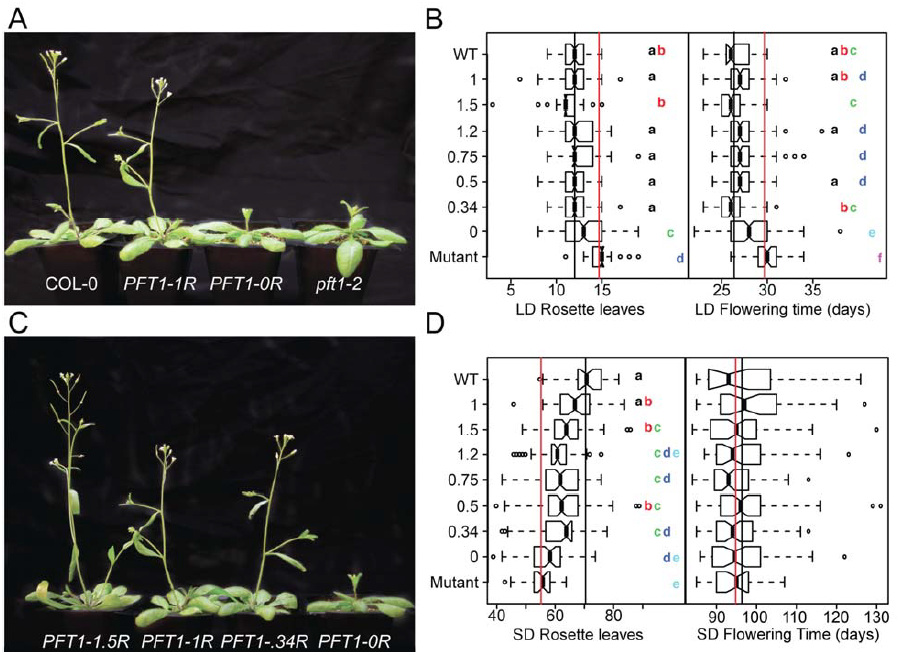
*PFT1* STR alleles differ in their ability to rescue a *pft1* loss-of-function mutant for flowering phenotypes. **A, C)** Transgenic plants carrying different *PFT1* STR alleles. Plants were grown under LD for 31 days and photographed. Background was removed in Adobe Photoshop CS 6.0. **B, D**) Strains sharing letters are not significantly different by Tukey’s HSD test. Black lines represent WT means, red lines represents *pft1-2* means for each phenotype. Each STR allele is represented by at least two independent transgenic lines (Table S3), with N > 20 for SD phenotypes and N > 35 for LD phenotypes, α = 0.05. LD = long days, SD = short days. In SD flowering time (days), no groups are significantly different.

In short days (SD), we observed an unexpected reversal in rosette leaf phenotypes (compare SD and LD rosette leaves, Figures 1B, 1D). Rather than flowering late (adding more leaves) as in LD, the loss-of-function *pft1-2* mutant appeared to flower early (fewer leaves at onset of flowering). Only the endogenous STR (1R) fully rescued this unexpected phenotype (Figure 1D). We observed the same mean trend for days to flowering in SD, although differences were not statistically significant, even for *pft1-2* (Figure 1D). This discrepancy may be due to insufficient power, or to a physiological decoupling of number of rosette leaves at flowering and days to flowering phenotypes in *pft1-2* under SD conditions. Regardless, our results indicate that *pft1-2*’s late-flowering phenotype is specific to LD conditions. Our observation of this reversal in flowering time-related phenotypes appears to contradict previous data (Cerdán and Chory 2003). However, a closer examination of this data reveals that the previously reported rosette leaf numbers in SD for the *pft1-2* mutant show a similar trend. *PFT1* STR length shows an approximately linear positive relationship with the SD rosette leaf phenotype, forming an allelic series of phenotypic severity. This allelic series strongly supports our observation of either slower growth rate (*i.e.* delayed addition of leaves) or early flowering of *pft1-2* as measured by SD rosette leaves at flowering.

*PFT1* genetically interacts with the red/far-red light receptor phyB, which governs petiole length through the shade avoidance response (Cerdán and Chory 2003; Wollenberg *et al*. 2008). We measured petiole length at bolting for plants grown under LD to evaluate the strength of their shade avoidance response, and thus whether the genetic interaction is affected by repeat length. Like the flowering time phenotypes, we found that the 1R allele most effectively rescued the long-petiole phenotype of the *pft1-2* null among all STR alleles (Figure 2), though some alleles (*e.g.* 1.5R) show a rescue that is nearly as good.

**Figure 2.**
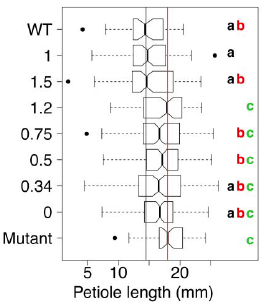
*PFT1* STR alleles differ in their ability to rescue a *pft1* loss-of-function mutant for petiole length in long days. Strains sharing letters are not significantly different by ANOVA with Tukey’s HSD test. Black lines represent WT means, red lines represent *pft1-2* means for each phenotype. Each allele is represented by at least two independent transgenic lines, N > 35 for each allele, α = 0.05.

In summary, plants expressing the 1R transgene most closely resembled wild-type plants across a range of adult phenotypes. In contrast, the other STR alleles showed inconsistent performance across these phenotypes, rescuing only some phenotypes or at times out-performing wild-type.

### *PFT1* STR alleles fail to rescue early seedling phenotypes

We next assessed quantitative phenotypes in early seedling development, some of which had been previously connected to *PFT1* function. Specifically, we measured hypocotyl and root length of dark-grown seedlings and examined germination in the presence of salt (known to be defective in *pft1* mutants) (Elfving *et al*. 2011). The *pft1-2* mutant showed the previously reported effect on hypocotyl length as well as a novel defect in root length (Figure 3A). None of the transgenic lines, including the one containing the 1R allele, effectively rescued these *pft1-2* phenotypes (Figure 3A). Similarly, 1R was not able to rescue the germination defect of *pft1-2* on high-salt media. However, both the 1.5R and 0.5R alleles were able to rescue this phenotype (Figure 3B). In summary, no single STR allele, including the endogenous 1R, was consistently able to rescue the early seedling phenotypes of the *pft1-2* mutant. One explanation for the failure of the endogenous STR (PFT1-1R) to rescue early seedling phenotypes is that the *PFT1* transgene represents only the larger of two splice forms. The smaller *PFT1* splice form, which we did not test, may play a more important role in early seedling development. To explore this hypothesis, we measured mRNA levels of the two splice forms in pooled 7-day seedlings grown under the tested conditions and various adult tissues at flowering in Col-0 plants. However, we found that both splice forms were expressed in all samples, and in all samples the larger splice form was the predominant form (data not shown). The possibility remains that downstream regulation or tissue-specific expression may lead to a requirement for the smaller splice form in early seedlings.

**Figure 3.**
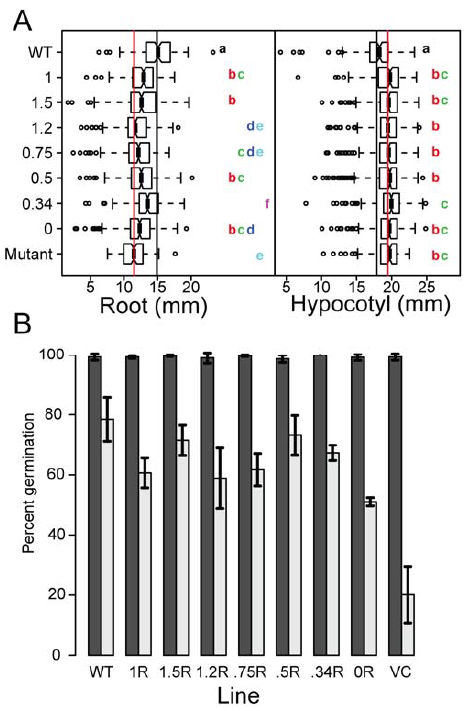
*PFT1* STR alleles differ in their ability to rescue a *pft1* loss-of-function mutant for early seedling phenotypes. **A)** Strains sharing letters are not significantly different by ANOVA with Tukey’s HSD test. Black lines represent WT means, red lines represent *pft1-2* means for each phenotype. Each allele is represented by at least two independent transgenic lines, N > 100 for all phenotypes for each allele, pooled across at least two experiments; α = 0.05. Hypocotyl length and root length were assayed in 7d seedlings grown in dark conditions. **B)** Dark and light bars represent mean germination across 3 biological replicates on 0 mM NaCl and 200 mM NaCl, respectively. N = 36 for each replicate experiment. Error bars represent standard error across these three replicates.

### Summarizing *PFT1* STR function across all tested phenotypes

Given the complex phenotypic responses to *PFT1* STR substitutions, results were equivocal as to which STR allele demonstrated the most ‘wild-type-like’ phenotype across traits, as measured by its sufficiency in rescuing *pft1-2* null phenotypes. To summarize the various phenotypes, we calculated the mean of each quantitative phenotype for each allele, and used principal component analysis (PCA) to visualize the joint distribution of phenotypes observed.

All STR alleles were distributed between the *pft1-2* null and wild-type (WT) in PC1, which was strongly associated with adult traits and represented a majority of phenotypic variation among lines (Figure 4). PC1 showed that 1R was the most generally efficacious allele for adult phenotypes. However, 1R showed incomplete rescue in early seedling phenotypes such as hypocotyl length, which drove PC2. All STR alleles showed substantial rescue in adult phenotypes, and even the 0R allele without an STR showed some partial rescue in some phenotypes; however, rescue of early seedling phenotypes was generally poor for all alleles. The first principal component also captured our observation that the *pft1-2* flowering defect reversed sign in SD vs. LD: according to Figure 4, SD and LD quantitative phenotypes are both strongly represented on principal component 1, but they show opposite directionality. We take this observation as support of this hitherto-unknown complexity in *PFT1* function.

**Figure 4.**
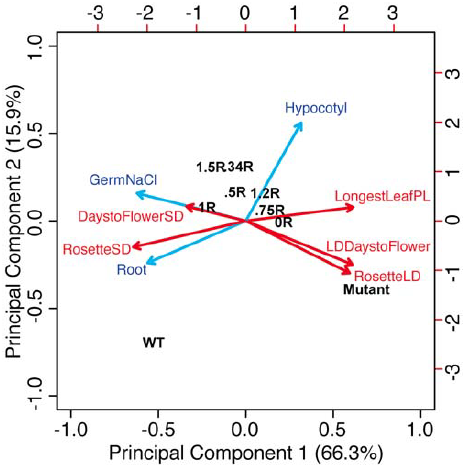
Distribution of *PFT1* STR allele performance across all phenotypes, relative to wild-type and *pft1-2* mutants. Biplot representation of PCA on all phenotypes across all tested *PFT1* STR alleles. Percentages on axes are the % variance in the overall data contributed by that principal component. Contributions of specific phenotypes to these axes are shown by size and direction of arrows. Red arrows represent adult phenotypes, blue arrows represent early seedling phenotypes; adult phenotypes are in red, whereas early seedling phenotypes are in blue. “RosetteSD”: number of rosette leaves under SD, “RosetteLD”: number of rosette leaves under LD, “LongestLeafPL”: petiole length of the longest leaf of rosette, “GermNaCl”: proportion of germinants on 200 mM NaCl, “Hypocotyl” and “Root” refer to lengths of the specified organs in dark-grown seedlings. Transgenic STR alleles are indicated by their proportion of the wild-type (WT) repeat, i.e. “1.5R”. Top and right axes provide a relative scale for the magnitude of phenotype vectors (blue and red arrows).

## DISCUSSION

STR-containing proteins pose an intriguing puzzle –they are prone to in-frame mutations, which in many instances lead to dramatic phenotypic changes (Gemayel *et al*. 2010). Although STR-dependent variation has been linked to adaptation in a few cases, the presence of mutationally labile STRs in functionally important core components of cell biology seems counterintuitive. PFT1, also known as MED25, is a core component of the transcriptional machinery across eukaryotes and contains an STR that is predicted to be highly variable in length. Contrary to this prediction, we found *PFT1* STR variation to be minimal, consistent with substantial functional constraint. The existing residual variation (∼2% of reference STR length, as opposed to > 100% for the *ELF3* STR in the same *A. thaliana* strains) suggests that the *PFT1* STR is mutationally labile like other STRs. In fact, several of the synthetic *PFT1* alleles examined in this study arose spontaneously during cloning. Strong functional constraint, however, may select against such deviations in STR length *in planta*.

Here, we establish the essentiality of the full-length *PFT1* STR and its encoded polyQ tract for proper PFT1 function in *A. thaliana*. We found that diverse developmental phenotypes were altered by the substitution of alternative STR lengths for the endogenous length. Leveraging the support of the *PFT1* STR allelic series, we report new aspects of PFT1 function in flowering time and root development.

### The *PFT1* STR is required for PFT1 function in adult traits

The *PFT1* 0R lines did not effectively complement *pft1-2* for adult phenotypes, suggesting a crucial role of the *PFT1* STR in regulating the onset of flowering and shade avoidance. Generally, *PFT1*-1R was most effective in producing wild-type-like adult phenotypes. The precise length of the STR, however, seemed less important for the onset of flowering in LD. With exception of *PFT1*-0R, all other STR alleles were also able to rescue the loss-of-function mutant to some extent, suggesting that as long as some repeat sequence is present, the PFT1 gene product can fulfill this function. Under other conditions, and for other adult phenotypes, requirements for *PFT1* STR length appeared more stringent. Specifically, under SD, the rosette leaf number phenotype of the *pft1-2* mutant can only be rescued by *PFT1*-1R, while STR alleles perform worse with increasing distance from this length “optimum”.

### *pft1-2* mutants are late-flowering in LD but not SD

*pft1-2* plants had fewer rosette leaves at flowering in SD, but more rosette leaves in LD, consistent with previous, largely undiscussed observations (Cerdán and Chory 2003). Under LD conditions, *pft1-2* null mutants flowered late, as described in several previous studies (Cerdán and Chory 2003; Wollenberg *et al*. 2008), but we observe no such phenotype under SD conditions, contradicting at least one prior study (Cerdán and Chory 2003). These data suggest that while PFT1 functions as a flowering activator under LD, its role is more complex under SD.

One recent study showed that PFT1 function in LD is dependent upon its ability to bind E3 ubiquitin ligases (Iñigo, Giraldez, *et al*. 2012). Inhibition of proteasome activity also prevents PFT1 from promoting *FT* transcription and thus inducing flowering, suggesting that degradation of PFT1 or associated proteins is a critical feature of *PFT1*’s transcriptional activation of flowering in LD. If this degradation is somehow down-regulated in SD, PFT1 could switch from a flowering activator to a repressor, through decreased Mediator complex turnover at promoters. Recent studies raised the possibility that different PFT1-dependent signaling cascades have different requirements for PFT1 turnover (Ou *et al*. 2011; Kidd *et al*. 2009), which may contribute to the condition-specific PFT1 flowering phenotype we observe. Conservatively, we conclude that the regulatory process that mediates the phenotypic reversal between LD and SD depends on the endogenous *PFT1* STR allele, suggesting that the polyQ is crucial to PFT1’s activity as both activator and potentially as a repressor of flowering.

### Incomplete complementation of germination and hypocotyl length by the PFT1 constructs

Whereas *pft1-2* adult phenotypes were rescued by the *PFT1*-1R allele, most of our transgenic lines could not fully rescue *pft1-2* early seedling phenotypes of 1) germination under salt, 2) hypocotyl length, and 3) root length. The *PFT1* gene is predicted to have two different splice forms, the larger of which was used to generate our constructs (both splice forms contain the STR). Several studies have shown that, under stress conditions, different splice forms of the same gene can play distinct roles (Yan *et al*. 2012; Leviatan *et al*. 2013; Staiger and Brown 2013). We note that the conditions under which *PFT1-*1R fails to complement are also potentially stressful conditions (artificial media, sucrose, high salt, dark). The shorter splice form of PFT1 may be required in signaling pathways triggered under stress conditions. We presume that the failure to complement results from a deficiency related to this missing splice form. However, hypocotyl length was the only trait in which all examined STR alleles resembled the *pft1-2* mutant. The significant functional differentiation among the STR alleles for root length and germination suggests that the large splice form does retain at least some function in early seedling traits.

### Implications for STR and *PFT1* biology

Coding and regulatory STRs have been previously studied and discussed as a means of facilitating evolutionary innovation (Verstrepen *et al*. 2005). However, this means of innovation is based upon the same sequence characteristics that promote protein-protein and protein-DNA binding (Escher *et al*. 2000; Schaefer *et al*. 2012), such that STR variability must be balanced against functional constraints. This balance has recently been described for a set of 18 coding dinucleotide STRs in humans, which are maintained by natural selection even though any mutation is likely to cause frame-shift mutations (Haasl and Payseur 2014). These results, coupled with our observations, lend credence to these authors’ previous argument that not all STRs act as agents of adaptive change (Haasl and Payseur 2013). Considering again the possibility that more conserved coding tandem repeats have distinct functions from non-conserved tandem repeats (Schaper *et al*. 2014), we suggest that *PFT1* and *ELF3* can serve as models for these two selective regimes, and that the structural roles of their respective polyQs underlie the differences in natural variation between the two. In some cases, such as *ELF3*, high variability is not always inconsistent with function, even while holding genetic background constant (Undurraga *et al*. 2012). In *PFT1*, we have identified a STR whose low variability reflects strong functional constraints. We speculate that these constraints are associated with a structural role for the PFT1 polyQ in the Mediator complex, either in protein-protein interactions with other subunits or in protein-DNA interactions with target promoters. Given that a glutamine-rich C-terminus appears to be a conserved feature of MED25 even in metazoans (File S1), we expect that our results are generalizable to Mediator function wherever this protein is present. Future work will be necessary in understanding possible mechanisms by which the MED25 polyQ might facilitate Mediator complex function and contribute to ontogeny throughout life. Moreover, attempts must be made to understand the biological and structural characteristics unique to polyQ-containing proteins that tolerate (or encourage) polyQ variation, as opposed to those polyQ-containing proteins (like PFT1) that are under strong functional constraints.

## METHODS

### Cloning

A 1000 bp region directly upstream of the *PFT1* coding region was amplified and cloned into the pBGW gateway vector (Karimi *et al*. 2002) to create the entry vector pBGW-*PFT1*p. A full-length *PFT1* cDNA clone, BX816858, was obtained from the French Plant Genomic Resources Center (INRA, CNRGV), and used as the starting material for all our constructs. The *PFT1* gene was cloned into the pENTR4 gateway vector (Invitrogen) and the repeat region was modified by site-directed mutagenesis with QuikChange (Agilent Technologies), followed by restriction digestions and ligations. The modified *PFT1* alleles were finally transferred to the pBGW-*PFT1*p vector via recombination using LR clonase (Invitrogen) to yield the final expression vectors. Seven constructs expressing various polyQ lengths (Table S2), plus an empty vector control, were used to transform homozygous *pft1-2* mutants by the floral dip method (Clough and Bent 1998). Putative transgenics were selected for herbicide resistance with Basta (Liberty herbicide; Bayer Crop Science) and the presence of the transgene was confirmed by PCR analysis. Homozygous T3 and T4 plants with relative *PFT1* expression levels between 0.5 and 4 times the expression of Col-0 were utilized for all experiments described. A minimum of two independent lines per construct was used for all experiments.

### Expression Analysis

All protocols were performed according to manufacture’s recommendations unless otherwise noted. Total RNA was extracted from 30 mg of 10-days-old seedlings with the Promega SV Total RNA Isolation System (Promega). 2 μg of total RNA were subjected to an exhaustive DNaseI treatment using the Ambion Turbo DNA-free Kit (Life Technologies). cDNA was synthesized from 100-300 ng of DNase-treated RNA samples with the Roche Transcriptor First Strand cDNA Synthesis Kit (Roche). Quantitative Real-Time PCR was performed in a LightCyler® 480 system (Roche) using the 480 DNA SYBR Green I Master kit. Three technical replicates were done for each sample. RT-PCR was performed under the following conditions: 5 min at 95 °C, followed by 35 cycles of 15 s at 95 °C, 20 s at 55 °C, and 20s at 72 °C. After amplification, a melting-curve analysis was performed. Expression of *UBC21* (At5g25760) was measured as a reference in each sample, and used to calculate relative *PFT1* expression. All expression values were normalized relative to WT expression, which was always set to 1.0. To measure splice forms, the protocol was the same but reactions were carried out in a standard thermal cycler and visualized on 2% agarose stained with ethidium bromide. For primers, see Table S4.

### Plant Materials and Growth Conditions

Homozygous plants for the T-DNA insertional mutant SALK_129555, *pft1-2*, were isolated by PCR analysis from an F2 population obtained from the Arabidopsis Stock Center (ABRC) (Alonso *et al*. 2003). Plants were genotyped with the T-DNA specific primer LBb1 (http://signal.salk.edu/tdna_FAQs.html) and gene-specific primers (Table S4).

Seeds were stratified at 4°C for 3 days prior to shifting to the designated growth conditions, with the shift day considered day 0. For flowering time experiments, plants were seeded using a randomized design with 15-20 replicates per line in 4×9 pot trays. Trays were rotated 180° and one position clockwise everyday in order to further reduce any possible position effect. Plants for LD were grown in 16 hours of light and 8 hours of darkness per 24 hour period. Bolting was called once the stem reached 1 cm in height.

Full strength MS media containing MES, vitamins, 1% sucrose, and 0.24% phytagar was used for hypocotyl experiments. For germination experiments, half-strength MS media was used, supplemented with 1% sucrose, 0.5 g/L MES, and 2.4 g/L phytagel containing 200 mM NaCl or H2O mock treatment with the pH adjusted to 5.7. All media was sterilized by autoclaving with 30 minutes of sterilization time. Seeds for tissue culture were surface sterilized with ethanol treatment prior to plating and left at 4°C for 3 days prior to shifting to the designated growth conditions. Plants for hypocotyl experiments were grown with 16 hours at 22°C and 8 hours at 20°C in continuous darkness following an initial 2 hour exposure to light in order to induce germination. Germination experiments were scored on day 4 under LD at 20-22°C. ImageJ software was utilized to make all hypocotyl and root length measurements. Raw phenotypic data are included as File S3.

### Statistical Analysis

All statistical analyses and plots were performed in R version 2.15.1 with α = 0.05 (R Development Core Team 2012). Phenotypic data were analyzed using the analysis of variance (ANOVA), followed by Tukey’s HSD tests for the differences of groups within the ANOVA. Tukey’s HSD is a standard post-hoc test for multiple comparisons of the means of groups with homogeneous variance that corrects for the number of comparisons performed. Principal component analysis was performed using the *prcomp()* function after scaling each phenotypic variable to mean = 0 and variance = 1 across lines (phenotypes are not measured on the same quantitative scale; for example, SD flowering time ranges from 80 to 140 days, whereas LD rosette leaves ranges ∼5-15 leaves).

### Sequence Analysis

Length of *ELF3* and *PFT1* STRs were determined by Sanger (dideoxy) sequencing. Raw sequencing data are included as File S2. PFT1 and MED25 reference amino acid sequences were obtained from KEGG (Ogata *et al*. 1999) and aligned with Clustal Omega v1.0.3 with default options (Sievers *et al*. 2011).

## ACKNOWLEDGMENTS

We are grateful to members of the Queitsch lab for valuable discussions. This work was supported by National Human Genome Research Institute Interdisciplinary Training in Genome Sciences Grants (2T32HG35-16 to M.O.P. and T32HG000035-16 to S.F.U.) and the Herschel and Caryl Roman Undergraduate Scholarship Fund (to J.B). The authors would like to thank the NIH/NHGRI Genome Training Grant, the National Institute of Health New Innovator Award (DP2OD008371 to C.Q.) and the Royalty Research Fund (RRF4365 to C.Q.) for their generous financial support.

## AUTHOR CONTRIBUTIONS

C.Q. and S.U. designed the research. P.R., J.B., T.G., M.O.P., and S.U. performed research. J.B. generated the transgenic lines. M.O.P., J.B. and S.U. analyzed data. S.U., M.O.P., P.R., J.B. and C.Q. wrote the paper.

